# The Mechanical and Biological Evolution of Pressure Ulcer Formation and Healing in Mice

**DOI:** 10.64898/2026.07.25.740666

**Authors:** Chien-Yu Lin, Shreya Sreedhar, Matthew J. Lohr, Colton J. Kostelnik, Alberto Madariaga, Adrian B. Tepole, Manuel K. Rausch

## Abstract

Pressure ulcers arise from sustained mechanical loading that impairs perfusion and damages skin tissues, yet the coupled mechanical and biological mechanisms of their formation and healing remain poorly characterized. We addressed this gap using a mouse model in which dorsal skin underwent 72 hours of magnet-induced ischemia followed by reperfusion, with tissue collected at 0, 3, 6, and 9 days and compared with baseline controls. From each mouse, we obtained paired samples from pressure ulcer and remote control (non-loaded) sites, mapped thickness by tissue profilometry, and performed equibiaxial testing with full-field digital image correlation and inverse finite element analysis to estimate regional material parameters. In parallel, we quantified CD31^+^ vasculature, F4/80^+^ macrophages, collagen content, and key cytokines. Pressure ulcer sites were compressed and thinner at Day 0, developed ulcers by Day 3, and continued to remodel through Day 9. Mechanical tests revealed heterogeneous strain fields with elevated deformation along ulcer borders, while remote control tissue deformed more homogeneously. These mechanical changes evolved alongside dynamic vessel and macrophage repopulation, increased collagen content at early time points, and cytokine upregulation within pressure ulcer tissue. Collectively, our data define the spatiotemporal co-evolution of tissue geometry, mechanics, collagen remodeling, and inflammation in pressure ulcers and provide a quantitative foundation for predictive mechanobiological models.

## Introduction

Pressure ulcers are a common and debilitating skin injury. In the US alone, pressure ulcers affect an estimated 1–3 million people annually [38]. In addition to causing discomfort and lowering quality of life, they are also a common cause of infection, thus increasing morbidity [52]. As a result, they contribute to the substantial cost of health care in the US and other health systems worldwide [40, 8]. Pressure ulcers form after sustained local compression that can occlude/distort microvasculature, leading to ischemia and, following pressure release, reperfusion injury [37, 5]. Risk is highest in individuals with immobility or inactivity, or sensory loss, who cannot reliably reposition or perceive injurious loading [38, 50]. Toward this end, a complete understanding of pressure ulcer formation and healing is pressing.

Sustained compression-induced ischemia followed by reperfusion initiates a coordinated yet dysregulated cellular and molecular response during pressure ulcer formation [47, 44, 39]. Early after hypoxia, resident skin cells upregulate chemotactic signals such as monocyte chemoattractant protein-1 (MCP-1). In cutaneous ischemia-reperfusion models, MCP-1 elevation precedes inflammatory cell infiltration, including macrophages commonly quantified as F4/80^+^ cells and neutrophils assessed through myeloperoxidase (MPO) activity [44, 51, 12, 18]. These recruited leukocytes amplify local tissue injury through pro-inflammatory mediators such as tumor necrosis factor-*α* (TNF-*α*) and interleukin-1*β* (IL-1*β*), which increase within the wound area in experimental pressure ulcer models [44, 49]. As healing progresses, macrophages contribute to tissue repair by coordinating inflammation resolution and producing growth factors. Studies that mitigate injury in pressure ulcer models report improved outcomes accompanied by increased expression of pro-repair mediators such as transforming growth factor-*β* (TGF-*β*), including TGF-*β*1, alongside preservation of vascular integrity [20, 51]. Consistent with this repair process, angiogenesis accompanies tissue restoration, with increased vascular endothelial growth factor (VEGF) and CD31^+^ endothelial cells indicating recovery of vascular integrity after ischemia–reperfusion injury [49].

Tissue injury, including cellular apoptosis and extracellular matrix damage, alters local tissue composition and structure, leading to dynamic changes in mechanical properties [10, 34, 45]. These alterations in stiffness and geometry may influence tissue deformation, microvascular perfusion, and susceptibility to continued ischemic injury, thereby potentially perpetuating pressure ulcer progression. Interestingly, while we have a descriptive understanding of the biology of pressure ulcer formation, we lack a complete mechanistic understanding [11]. Moreover, the mechanical evolution of ulcerated skin remains poorly characterized, and, to our knowledge, no prior studies have quantified its temporal mechanical evolution in vivo. That is, the interplay between biology and tissue mechanics in a longitudinal setting lacks quantitative data and, thus, poses an obstacle to our understanding. Moreover, there is a near-complete lack of multi-scale and multi-modal data that includes tissue-level, cell-level, and gene-level markers of damage and healing.

Recognizing the need for a quantitative understanding of these coupled processes, our prior work has focused on developing computational models of pressure ulcer formation and healing. Given the complex interplay between tissue mechanics and biology, we developed a multi-physics modeling framework that integrates a systems-biological approach with a mechanics solver to account for tissue strain and stress, as well as the mechanobiological coupling of both [47, 46]. However, limited experimental data are available to inform and validate our or other similar models [54, 31, 17]. Therefore, we miss out on the opportunity to build tools that could serve pressure ulcer diagnosis, prognoses, or function as a virtual test bed for novel therapies.

The goal of this current work is to fill these gaps. That is, we set out to collect longitudinal data on the mechanical and biological co-evolution of pressure ulcers. We do so by characterizing the mechanics and geometry of injured and healing skin, the evolution of its structural and cellular composition, and the accompanying molecular responses. To this end, we captured the longitudinal evolution of tissue structure (thickness maps and histology), mechanics (biaxial stiffness), cellular responses (CD31^+^ and F4/80^+^ cells), inflammatory signaling (MPO, TNF-*α*, MCP-1, IL-1*β*, VEGF, and TGF-*β*1), extracellular matrix composition (collagen), and gene expression (RNA-seq) in a well-established mouse model. Through our work, we provide critical data for our own and others’ efforts toward developing quantitative models of pressure ulcers and their healing.

## Methods and Materials

### Pressure Ulcer Mouse Model

In our work, we used 13-to 15-week-old male C57BL/6 mice. In all experiments, we strictly followed the NIH Guide for the Care and Use of Laboratory Animals, and the Institutional Animal Care and Use Committee at The University of Texas at Austin approved all procedures under #AUP-2023-00092. We anesthetized mice with 2% isoflurane in oxygen at a flow rate of 2 L/min, delivered via a nose cone. While anesthetized, we removed dorsal hair using electric clippers followed by a chemical depilatory agent (Nair, Church & Dwight Co., Inc., Ewing, NJ, USA) to expose the skin for pressure application.

Building on the dorsal skin ischemia–reperfusion model described by Stadler et al., we applied a pair of cylindrical neodymium magnets (McMaster-Carr Part #5862K103) mounted in custom-machined plastic holders to the dorsal skin [48]. The magnets generated a sustained compressive pressure of approximately 150 mmHg over a circular contact area of 5/16 inch (7.94 mm) in diameter, which we maintained for 72 hours to induce ischemic injury. After magnet removal, we survived mice for 0-, 3-, 6-, and 9-days (Figure 1**a**).

**Figure 1:**
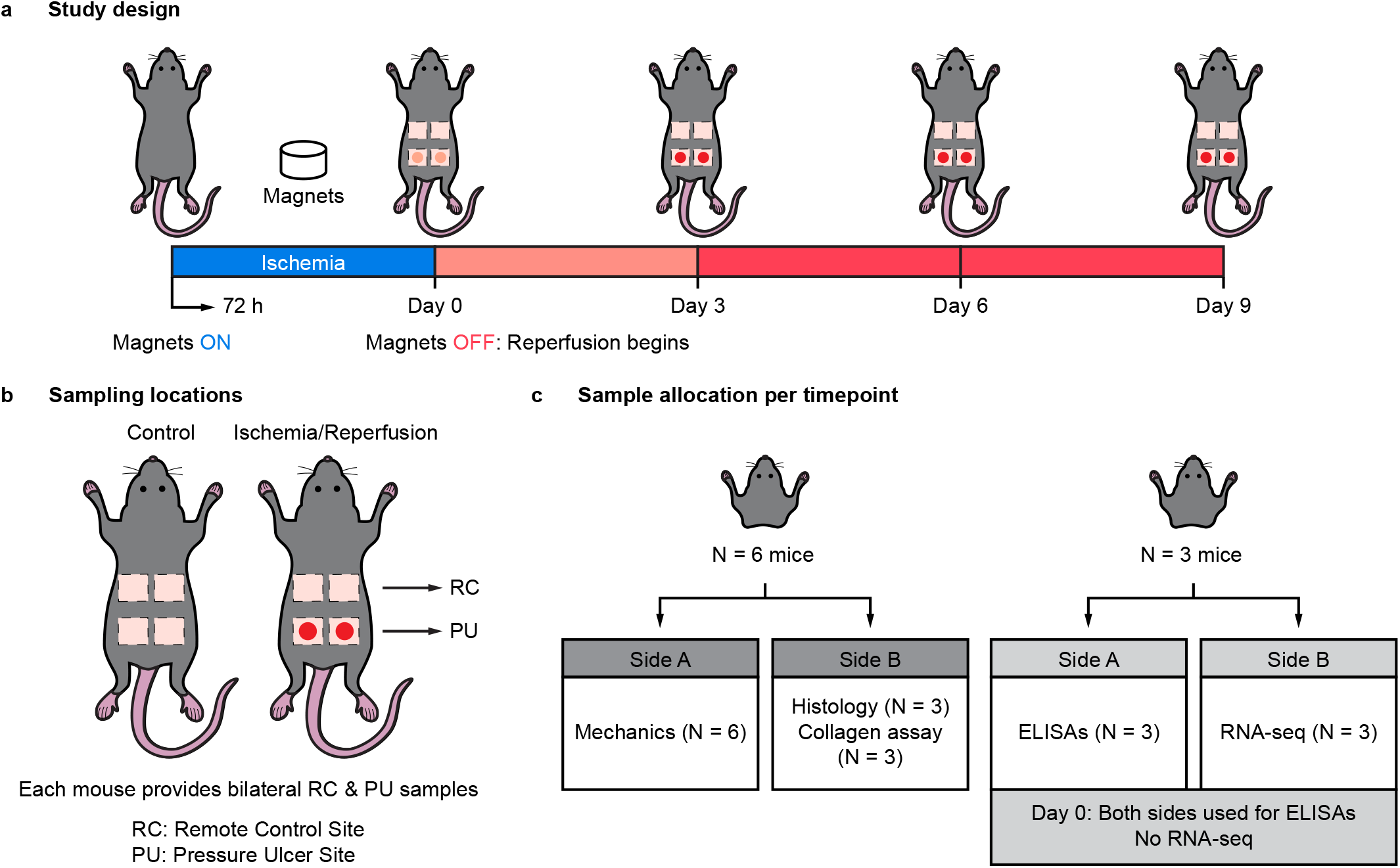
: Study design for the pressure ulcer model, including **a**) ischemia-reperfusion timeline, **b**) sampling locations (RC and PU), and **c**) tissue allocation across mechanical and biological assays

In each mouse, we collected samples from both the pressure ulcer (PU) sites and anatomically distant remote control (RC) regions (Figure 1**b**). Note, by the virtue of our model, each experiment yielded two ulcer samples per animal, one on each side of the dorsal midline. Age-matched mice without magnet application served as controls.

For biomechanical characterization, we used six mice per group (N = 6) and randomly selected skin samples from one side of each mouse. We allocated the contralateral side to histological staining (N = 3) and collagen content quantification (N = 3); see Figure 1**c**. In a separate cohort (N = 3 mice per group), we collected bilateral skin samples. For Control and Days 3, 6, and 9, we used one side for ELISA-based inflammatory cytokine analysis (N = 3) and the contralateral side for RNA-seq (N = 3). At Day 0, reduced tissue thickness resulted in insufficient sample mass for homogenization; therefore, we allocated both sides to ELISAs and did not perform RNA-seq at this time point. For collagen assays and ELISAs, we cryogenically stored 8 mm skin biopsy punches at -80 ^*o*^C in a 9:1 DMEM:DMSO solution supplemented with protease inhibitor (ThermoFisher, A32953, Waltham, MA) immediately after excision until downstream processing.

### Mechanical Testing

We excised 12 mm × 12 mm square skin samples for biomechanical characterization. Prior to biaxial testing, we obtained thickness profiles of each skin sample using our previously developed 3D profilometry imaging technique [26]. We then speckled the epidermal side with graphite powder for digital image correlation (DIC). After mounting the sample on our biaxial device (Biotester, Cellscale, Waterloo, ON, Canada) and submerging the sample in 1 × PBS at 37 ^*o*^C, we preloaded the tissue equibiaxially (the same in all directions) to 50 mN to establish a consistent reference state and remove tissue slack. Next, we performed 20 preconditioning cycles equibiaxially to 1000 mN and then conducted two final equibiaxial cycles to 1000 mN (force-controlled test at a quasi-static rake displacement rate of approximately 0.25 mm/s). We chose this force to induce large deformations in the skin samples without causing damage. While testing, we continuously captured images of the speckle pattern at 5 Hz for offline DIC.

We performed all tests within 4 hours after excision. We acquired actual tissue strain and displacement during biaxial testing via DIC of the recorded graphite pattern using the open-source 2D-DIC software Ncorr [2]. Consistent with our previous work [36], we used only the final loading cycle in our analysis because we were interested in the response of equilibrated tissue, and we chose the end of the second downstroke as the reference configuration for strain and displacement calculations. Ncorr computed the in-plane Green–Lagrange strain components. We then calculated the maximum principal Green–Lagrange strain (*E*_max_) at each point as the larger principal value of the two-dimensional strain tensor and generated full-field maps to visualize the spatial distribution of tissue deformation.

### Inverse Finite Element Analysis

To quantify the mechanical properties of skin with and without pressure ulcers under equibiaxial tension, we developed an inverse finite element pipeline. In the following analysis, we report model fits for force-displacement data and displacement fields in both directions of loading.

We created finite element models to fit data from all 59 samples (12 Control, 23 RC, 24 PU). To this end, we first digitized the chosen reference configuration of our samples using Coreform Cubit 2022.11 (Coreform LLC, Orem, UT). Based on a thorough mesh convergence study, we opted to use a mesh length of *h* = 0.15 mm for all samples. Using a custom MATLAB 2024b (Mathworks, Natick, MA) program, we created input files for Abaqus Standard (Abaqus/CAE 2024, Simulia) to simulate equibiaxial tension for all skin samples. We assigned 2D linear plane stress elements (CPS4) to the skin samples, and to each element, we assigned a numerical thickness as measured using profilometry. Next, we assigned actual displacements as measured with DIC via Ncorr to each boundary node. Because data at the boundary were noisy, we smoothed the boundary displacements using a moving window average with a window of 10 displacement points.

To capture the mechanical response of skin samples to equibiaxial tension, we assigned different hyperelastic material laws to the skin and ulcer regions of our samples. We assigned the anisotropic Holzapfel-Gasser-Ogden (HGO) model [9, 25] to the skin regions of the control and PU samples. We modeled skin using one fiber family, as done before, [36]. The HGO model with one fiber family takes the following form,

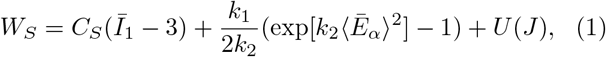

with *U* (*J*) defined as

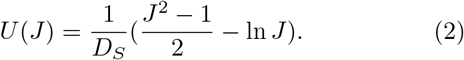

Here, *Ī*_1_ is the first invariant of the deviatoric right Cauchy-Green deformation tensor, 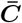, which is defined as 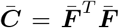 with 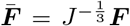 is the deformation gradient tensor and *J* = det[***F***]. Please note that here and throughout the text we will use upper case bold letters to refer to second-order tensors and lower case bold letters to refer to first-order vectors. The operator, ⟨•⟩, is the Macaulay bracket, which is defined as

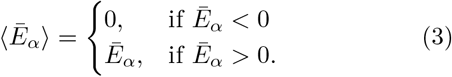

Here, 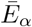 is defined as

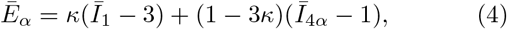

with 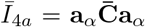. This term accounts for the strain energy of fibers, and the Macaulay brackets ensures that fibers have no compressive stiffness. The vector, **a**_*α*_, captures the mean orientation of the fibers based on the mean fiber angle, *γ* ∈ [0, *π*], relative to the X-axis. We let **a**_**1**_ = (cos[*γ*], sin[*γ*]). To capture the directionality of the HGO model, we assigned ***a***_**1**_ as the orientation vector for each skin element.

Because the underlying fibrous microstructure of the skin is damaged during the formation of pressure ulcers, as shown in [15], we assigned the Neo-Hookean model to the ulcer regions of PU samples. This model takes the form,

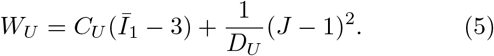

In the prior equations, _*S*_ represents skin-specific terms and _*U*_ represents ulcer-specific terms. The moduli *C*_*i*_ are related to the shear moduli, *µ*_*i*_, by 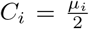, and *D*_*i*_ are related to the bulk moduli by 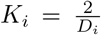 for *i* = (*S, U*). Note that we set 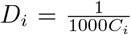 to achieve the near incompressible behavior exhibited by skin [27]. The parameter *k*_1_ captures the modulus of the fiber families; *k*_2_ defines the strain stiffening behavior of the fibers, and 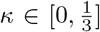 represents the dispersion of the fibers.

To identify the aforementioned parameters (*C*_*S*_, *C*_*U*_, *k*_1_, *k*_2_, *κ, γ*), we used a least-squares approach, whereby we minimized the error between the measured and predicted biaxial forces and displacement fields. For additional details regarding our approach, please see Supplemental Information 1.

### Histology & Immunohistochemistry

Upon excision, we immediately fixed 12 mm ×12 mm square skin samples allocated for histology in 10% neutral buffered formalin for 24 hours, then transferred them directly to 70% ethanol. A commercial histology service (Histoserv Inc., Germantown, MD, USA) prepared all histological slides by embedding them in paraffin, sectioning them longitudinally to a thickness of 5 *µ*m and staining them with hematoxylin and eosin (H&E), as well as immuno-histochemical staining for CD31 and F4/80 markers.

We acquired stitched histological images using a light microscope (Ti2-E High-Content Analysis System, Nikon, Tokyo, Japan) at 10× magnification. We used H&E staining to assess pressure ulcer progression by evaluating tissue morphology across experimental groups and time points. To enable spatial comparisons in immunohistochemistry (IHC) images, we divided each cross-sectional image into nine equal regions in a 3 × 3 grid along the lengthwise and thickness-wise axes. Using this regional segmentation, we quantified the percentage of positive staining for blood vessels (CD31^+^) and macrophages (F4/80^+^) with a custom MATLAB program [35, 22].

### Quantitative Collagen Assay

Immediately before testing, we rapidly thawed the samples to room temperature, rinsed with 1× PBS, dried them with Kimwipes, and recorded the wet mass of each sample. We added 100 *µ*L of deionized water for every 10 mg of wet tissue to homogenize the samples.

We hydrolyzed 100 *µ*L of homogenate in 100 *µ*L of 10 N NaOH at 120 °C for 1 h, then neutralized it with 100 *µ*L of 10 N HCl. After vortex mixing at 2000 ×g for 5 min, we transferred 10 *µ*L of hydrolysate to each well in triplicate and evaporated it to dryness on a 65 ^*o*^C heating plate. Following the Total Collagen Assay Kit protocol (BioVision Inc, K406, Milpitas, CA, USA), we measured colorimetric absorbance at 560 nm using a spectrophotometer (Tecan, Infinite 200 Pro, Männedorf, Switzerland) and interpolated collagen concentrations from a standard type-I linear fit curve (R2 = 0.99 ± 0.01).

### Inflammatory Marker Quantification

To prepare tissue lysates, we rapidly thawed the samples to room temperature and rinsed them with 1× PBS. We then dried the tissue samples with Kimwipes and weighed each one. We lysed skin samples in RIPA buffer (Sigma-Aldrich, R0278, St. Louis, MO, USA) supplemented with a protease inhibitor cocktail (Sigma-Aldrich, P8340, St. Louis, MO, USA), adding 10 *µ*L of protease inhibitor per 1 mL of RIPA buffer. We used 100 *µ*L of RIPA buffer with inhibitor per 10 mg of tissue. We placed the processed tissue in 2 mL microtubes with eight metal beads and added the RIPA buffer with inhibitor. We homogenized the tissue at full power for 30 seconds, placed the tubes on ice to cool, and repeated this cycle until no visible chunks remained. After homogenization, we centrifuged the samples at 4500 rpm for 10 minutes at 4 ^*o*^C. We collected the supernatant, aliquoted it for each assay, and stored it at -80 ^*o*^C until analysis.

We measured levels of the inflammatory markers myeloperoxidase (MPO; DY3667), monocyte chemoattractant protein-1 (MCP-1; DY479), transforming growth factor *β*1 (TGF-*β*1; DY1679), tumor necrosis factor *α* (TNF-*α*; DY410), interleukin 1*β* (IL1-*β*; DY401), and vascular endothelial growth factor (VEGF; DY493) using DuoSet ELISA kits (Bio-Techne, Minneapolis, MN, USA). We performed ELISAs according to manufacturer protocols and acquired absorbance data using a microplate reader. We calculated cytokine concentrations using standard curves provided with each kit. To normalize cytokine levels to total protein, we measured the protein concentration of each lysate using the Pierce™ Microplate BCA Protein Assay Kit (Thermo Fisher Scientific, 23252, Waltham, MA, USA). We ran all measurements in triplicate.

### RNA Sequencing

To preserve RNA integrity, we stored each sample in RNAlater (Invitrogen) at -20°C until RNA extraction. We extracted total RNA from all samples using the Qiagen RNeasy Fibrous Tissue Mini Kit (Qiagen, USA) and quantified RNA concentration and quality using Nanodrop spectrophotometry. Normalized RNA concentrations were then submitted to The University of Texas at Austin Genomic Sequencing and Analysis Facility for RNA integrity assessment, library preparation, and 3’ Tag Sequencing (TagSeq) [30]. Libraries were quantified using the Quantit PicoGreen dsDNA assay (ThermoFisher) and pooled equally for subsequent size selection at 350-550 bp on a 2% gel using the Blue Pippin (Sage Science). The final pools were checked for size and quality with the Bioanalyzer High Sensitivity DNA Kit (Agilent) and their concentrations were measured using the KAPA SYBR Fast qPCR kit (Roche). The samples were then sequenced on the NovaSeq 6000 (Illumina) instrument with single-end, 100-bp reads. TagSeq library deduplication and adapter trimming were performed by The University of Texas at Austin Computation Biology and Bioinformatics Core Facility (RRID:SCR 022688). All computational analyses were conducted on the Texas Advanced Computing Center (TACC) HPC cluster using Apptainer biocontainers. We aligned high-quality 3’ TagSeq reads to the *Mus musculus* reference genome (Mus musculus.GRCm39.cdna.all.fa) using STAR (v2.7.11b). We then assigned annotated gene features to aligned reads using HTSeq-count (v2.0.7). After the alignment and gene assignment steps, we used MultiQC to assess mapping quality. To analyze differential gene expression, we used R (v3.3.2) using a Biocontainer package, DESeq2. The differential gene expression model included the sample location (RC & PU) & group (Control, Day 3, 6 & 9) as fixed effects paired by animal. We normalized gene counts using regularized log transformation, and p-values were adjusted for multiple comparisons. Relevant genes were further analyzed using a linear mixed model that accounted for day and location as fixed effects and subject as a random effect. Genes with an adjusted p-value *<* 0.05 were considered significantly differentially expressed.

### Analysis and Statistics

We conducted all statistical analyses in R (Version 4.5.2), where statistical significance was defined as p *<* 0.05. For all data, we used a linear mixed model as implemented in the R package *afex* (Version 1.4.1). All posthoc analyses were conducted using Tukey post-hoc tests. Unless indicated otherwise, we report data as mean ±1 Standard Deviation.

## Results

### Pressure ulcers display temporally evolving thickness remodeling and strain concentration

We first characterized the temporal evolution of pressure ulcer structure and mechanical behavior following ischemia-reperfusion injury. Representative images (Figure 2**a**) showed the emergence and progression of wound morphology over time, beginning at Day 0 and becoming more pronounced through Days 3–9. To quantify these structural changes, thickness maps derived from tissue profilometry (Figure 2**b**) revealed pronounced spatial heterogeneity within pressure ulcer (PU) samples. At Day 0, ulcer regions exhibited localized thinning relative to surrounding tissue. Over subsequent time points, these regions displayed localized thickening, reflecting temporally evolving structural remodeling. Consistent with these geometric changes, maximum principal Green–Lagrange strain (*E*_max_) maps (Figure 2**c**) demonstrated non-uniform strain distributions across the tissue surface. Notably, elevated strain concentrations emerged at the periphery of ulcer regions beginning at Day 0 following reperfusion injury. These strain patterns evolved over time, reflecting progressive changes in tissue load-bearing behavior during ulcer progression and healing.

**Figure 2:**
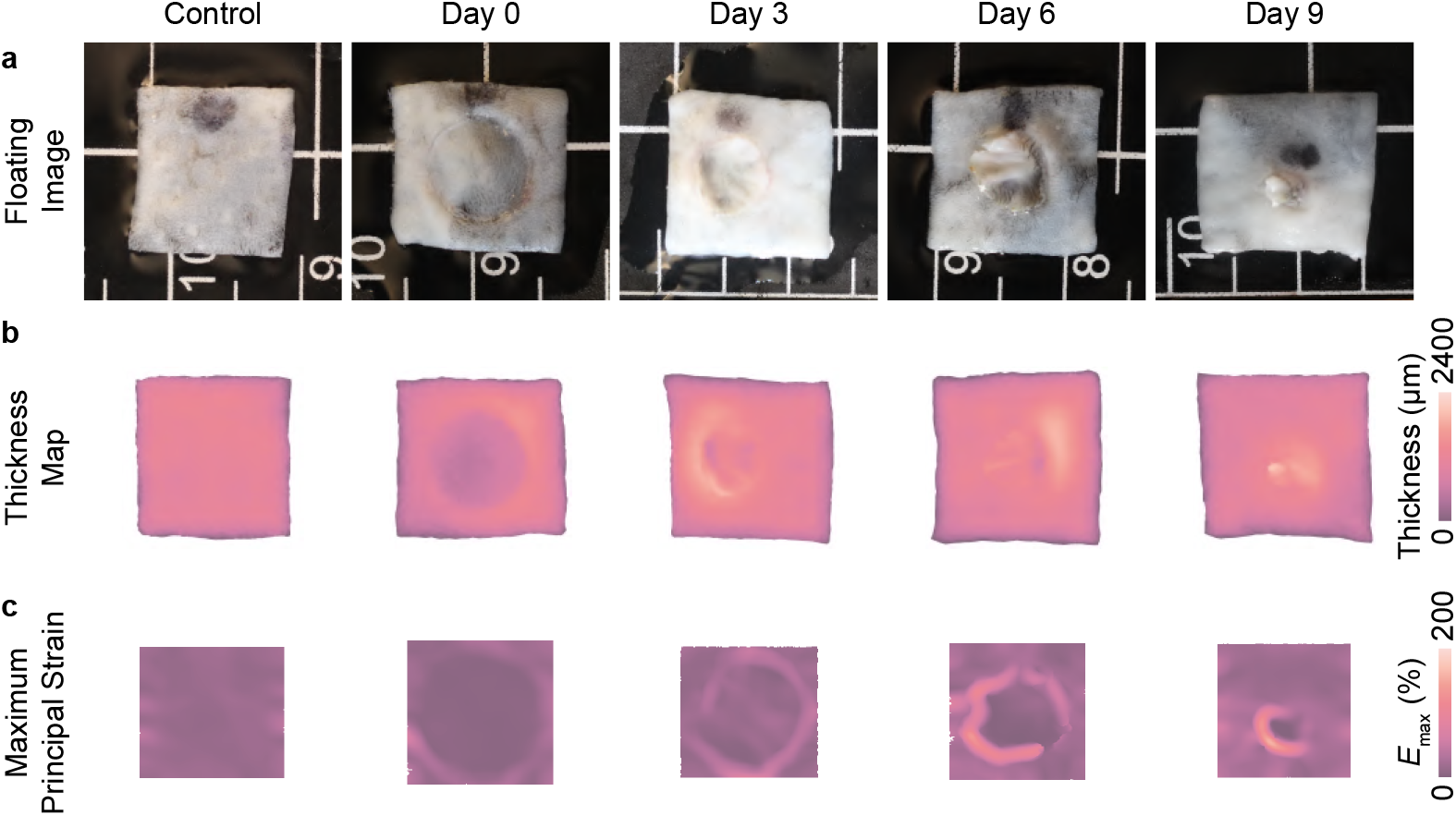
: Pressure ulcers display temporally evolving thickness remodeling and strain concentration. Representative **a**) images, **b**) thickness maps, and **c**) maximum principal Green–Lagrange strain (*E*_max_) maps show localized thinning/thickening and strain distribution on the pressure ulcer (PU) samples over time

### Pressure ulcer formation is associated with early increases in matrix moduli and fiber stiffening parameters

The experimental force and displacement data, as well as the displacement fields, were fit using the approach previously described. Exemplary fits to the force and displacement data for control and PU samples are shown in Figure 3. The average NMSE across all fits is 0.93, demonstrating that our modeling approach is suitable for capturing the mechanical properties of skin with and without pressure ulcers. Further details on the quality-of-fit metrics can be found in the Supplementary Table S1.

**Figure 3:**
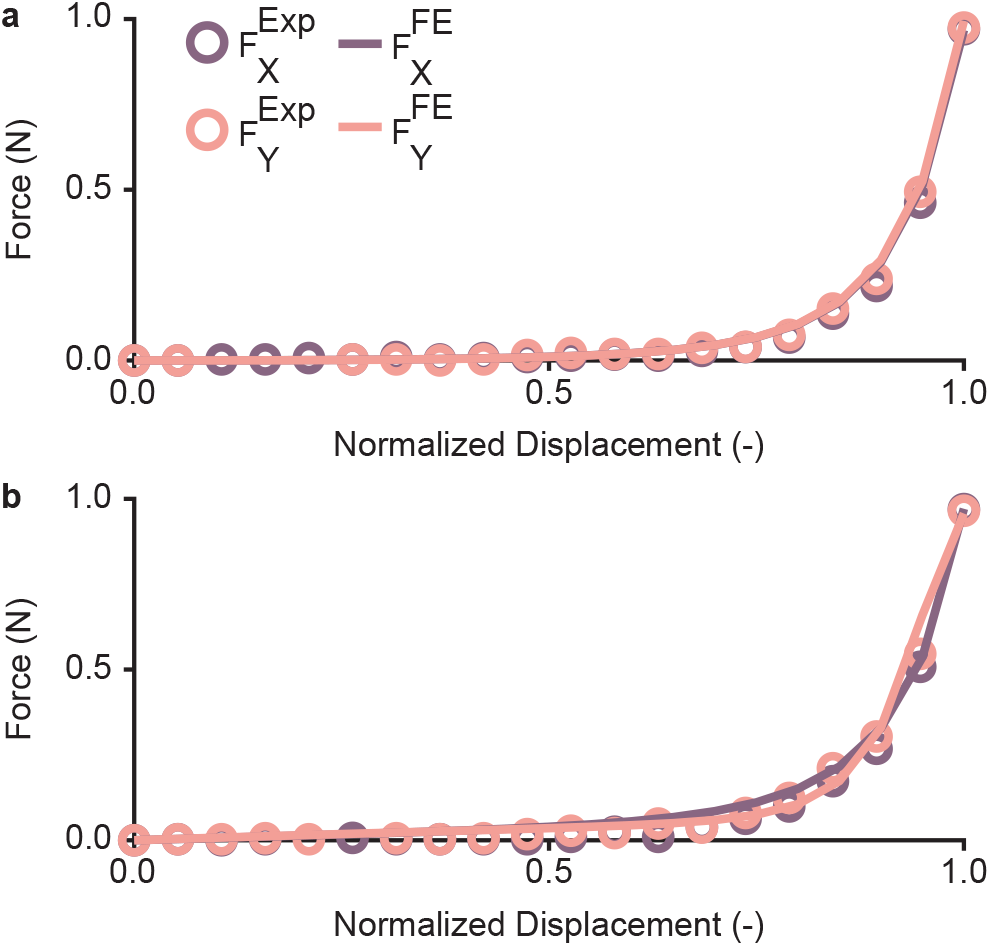
: Exemplary fits to X- and Y-force and displacement data for **a**) a control sample and **b**) a pressure ulcer (PU) sample. The normalized mean squared errors (NMSE) for the fits to the control sample are 0.99 for both the X- and Y-forces. The NMSEs for the fits to the PU sample are also 0.99 for both the X- and Y-forces

We show the parameters for all fits in Figure 4. Note that we only show parameters governing material stiffness: *C*_*S*_, *C*_*U*_, *k*_1_, and *k*_2_. We do not show the parameters governing fiber orientation and dispersion: *γ* and *κ*. The matrix moduli of the ulcers (PU-Ulcer) were higher than those of the surrounding skin (PU-Skin) on Day 0 and decreased as the ulcers healed (Figure 4**a**). The fiber moduli of the skin tissues immediately surrounding the ulcers (PU-Skin) decreased on Day 0, implying that the skin damage extends beyond the ulcer regions (Figure 4**b**). The fiber stiffening parameter *k*_2_ showed distinct dynamics in these skin regions: in remote control (RC) skin, *k*_2_ decreased at Day 0 relative to control and remained relatively low at later time points, whereas in PU-Skin, *k*_2_ was elevated at Day 0 but decreased by Day 3 and remained lower than control thereafter (Figure 4**c**). These mechanical parameters suggest that, under the loading conditions used in our biaxial tests, ulcer tissue exhibits increased matrix stiffness relative to the surrounding skin before substantial collagen fiber recruitment and that damage from the formation of a pressure ulcer extends to the microstructure beyond the ulcer’s boundary.

**Figure 4:**
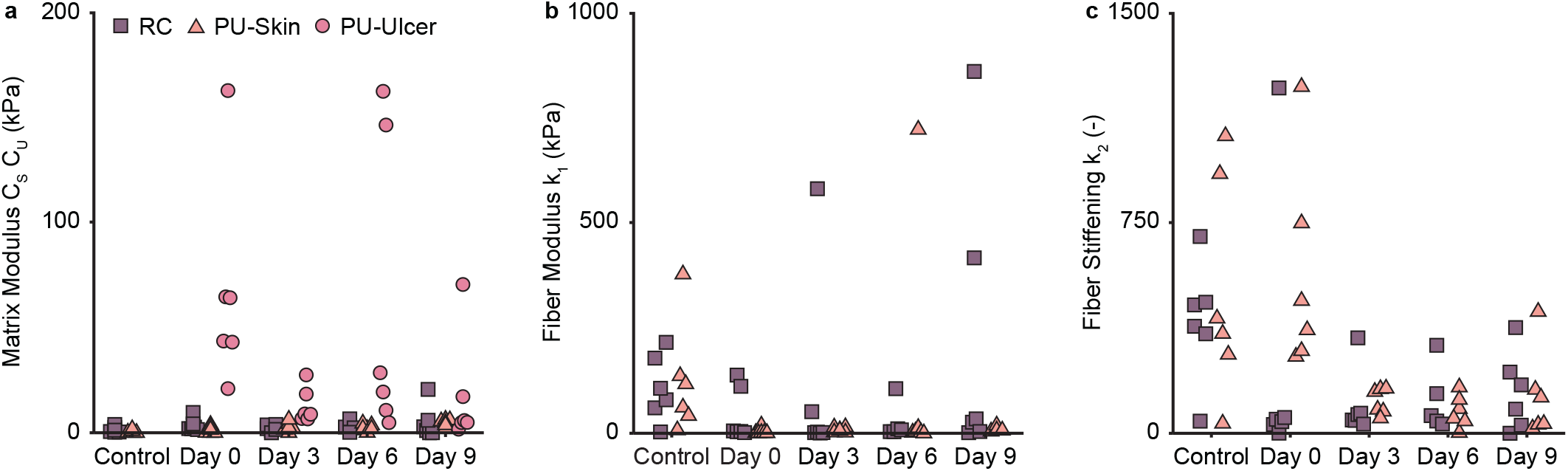
: Pressure ulcer formation is associated with early changes in matrix modulus and fiber mechanics. **a**) Matrix modulus *C*_*s*_ for the skin regions in remote control (RC) and pressure ulcer (PU) samples, and *C*_*u*_ for the ulcer region in the PU samples; pressure ulcers are initially stiffer than surrounding skin and soften over time. **b**) Fiber modulus *k*_1_ for the skin regions in both sample types; *k*_1_ decreases in PU samples, indicating that damage extends into skin surrounding the ulcer. **c**) Fiber stiffening parameter *k*_2_ for the skin regions in both sample types; in RC skin, *k*_2_ decreases at Day 0 and remains relatively low at later time points, whereas in PU-Skin regions *k*_2_ is elevated at Day 0 but decreases by Day 3 and remains lower than control thereafter, reflecting transient and sustained changes in strain-stiffening behavior. Note: RC = remote control samples, PU-Skin = skin region of PU samples, and PU-Ulcer = ulcer region of PU samples

### Pressure ulcer progression involves coordinated spatial re-organization of angiogenic and inflammatory responses

We provide representative H&E and IHC images from the PU site across all time points in Supplementary Figures S2-4. H&E staining revealed progressive tissue injury following ischemia-reperfusion, characterized by early disruption of dermal architecture and cellular organization at Day 0, followed by pressure ulcer formation and gradual tissue reorganization during healing. Figure 5**a** illustrates the regional segmentation scheme that we used for spatial quantification of the IHC images. Spatial quantification of CD31^+^ endothelial staining (Figure 5**b**) revealed early suppression within the ischemic center at Day 0 relative to control tissue. Following reperfusion, CD31^+^ signal increased predominantly in the side and deep regions at Days 3–6, suggesting progressive vascular recruitment toward previously ischemic areas. F4/80^+^ macrophage staining (Figure 5**c**) exhibited a similar spatiotemporal pattern. Macrophage density was reduced within the ulcer center at Day 0 but markedly elevated in the side and deep regions at Days 3–6. Notably, relative macrophage signal decreased in RC tissue at Day 3 compared to control mice, consistent with immune cell redistribution toward ulcer sites. By Day 9, macrophage distribution became more uniform across regions. Together, these findings demonstrate a coordinated, spatially structured inflammatory and angiogenic response characterized by early central suppression followed by peripheral-to-central recruitment during ulcer progression.

**Figure 5:**
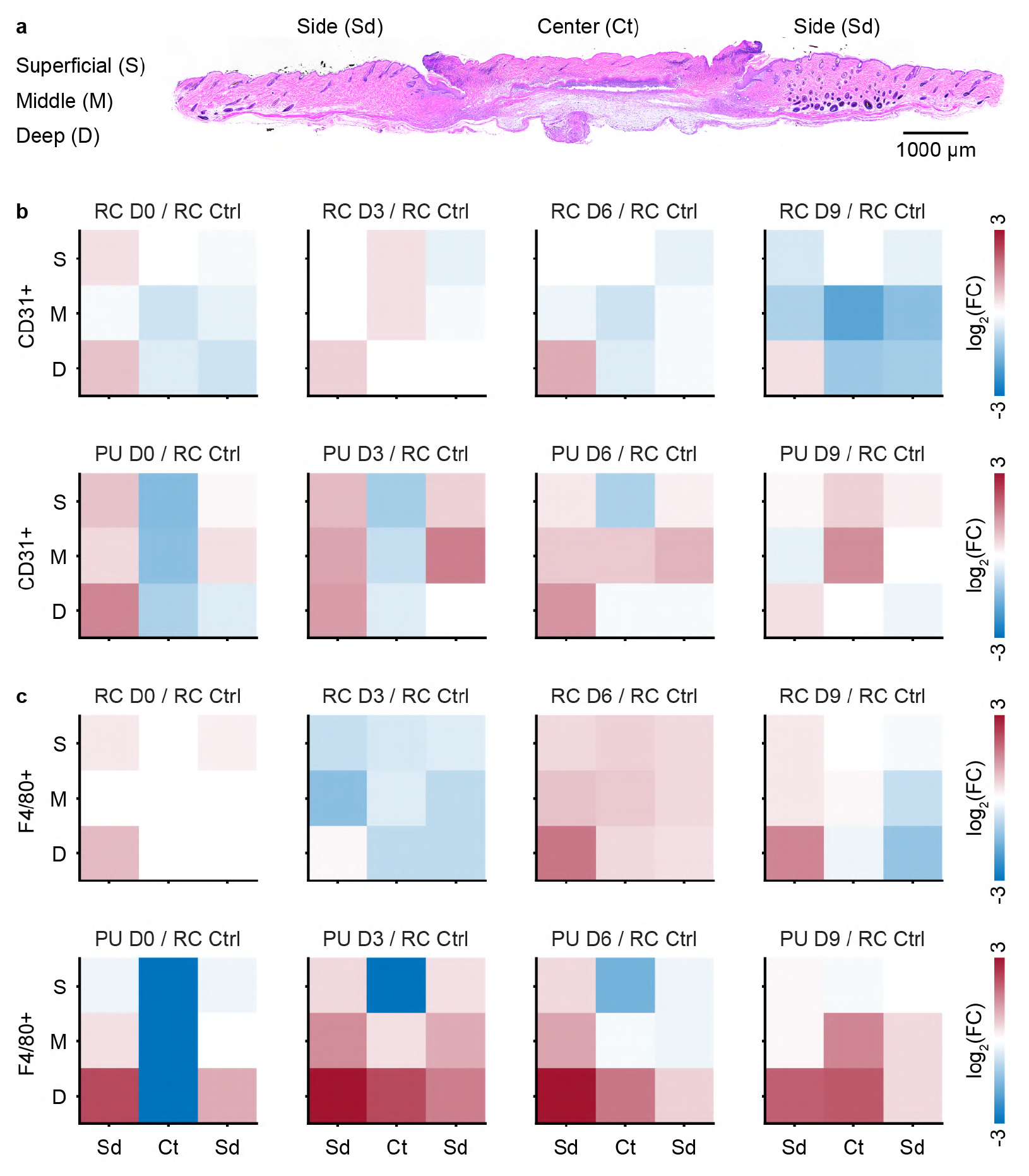
: Pressure ulcer progression involves coordinated spatial reorganization of angiogenic and inflammatory responses. **a**) Representative cross-sectional H&E image of a Day 6 pressure ulcer (PU) sample illustrating the regional segmentation scheme that we used for spatial analysis of the immunohistochemistry (IHC) images. We divided the ulcer center (Ct) and lateral sides (Sd) into superficial (S), middle (M), and deep (D) dermal layers for spatial quantification. Supplementary Figure S2 shows representative H&E images from the PU site across all time points, and Supplementary Figures S3–S4 show the corresponding CD31 and F4/80 IHC images. **b-c**) Heatmaps show regional log_2_ fold change of CD31^+^ endothelial cells and F4/80^+^ macrophages in PU samples relative to remote control (RC) samples in control mice across timepoints (N = 3)

### Pressure ulcer formation drives collagen remodeling at the protein and gene levels

Next, we evaluated collagen remodeling at both the protein and gene levels (Figure 6). We quantified total collagen content using a hydroxyproline assay (Figure 6**a**) and found that collagen content increased significantly at Day 0 in both PU and RC samples compared to control. Collagen content remained elevated at Day 3, Day 6, and Day 9, although levels gradually decreased at later time points. To assess gene expression, we analyzed RNA-seq data for major collagen genes associated with skin structure and function (Figure 6**b–f**). We also provide volcano plots showing upregulated and downregulated genes at each time point in Supplementary Figure S5. In PU samples, we observed increased expression of fibrillar collagen-encoding genes *Col1a1* and *Col3a1* at Days 3 and 6 relative to controls. We also detected time-dependent up-regulation of nonfibrillar collagen-encoding genes *Col4a2, Col6a1*, and *Col7a1*. In contrast, RC samples did not exhibit consistent upregulation of these genes. Collectively, these results show that collagen accumulation occurs early during ulcer progression, while sustained upregulation of collagen-encoding genes primarily occurs within ulcer tissue, supporting ongoing tissue remodeling.

**Figure 6:**
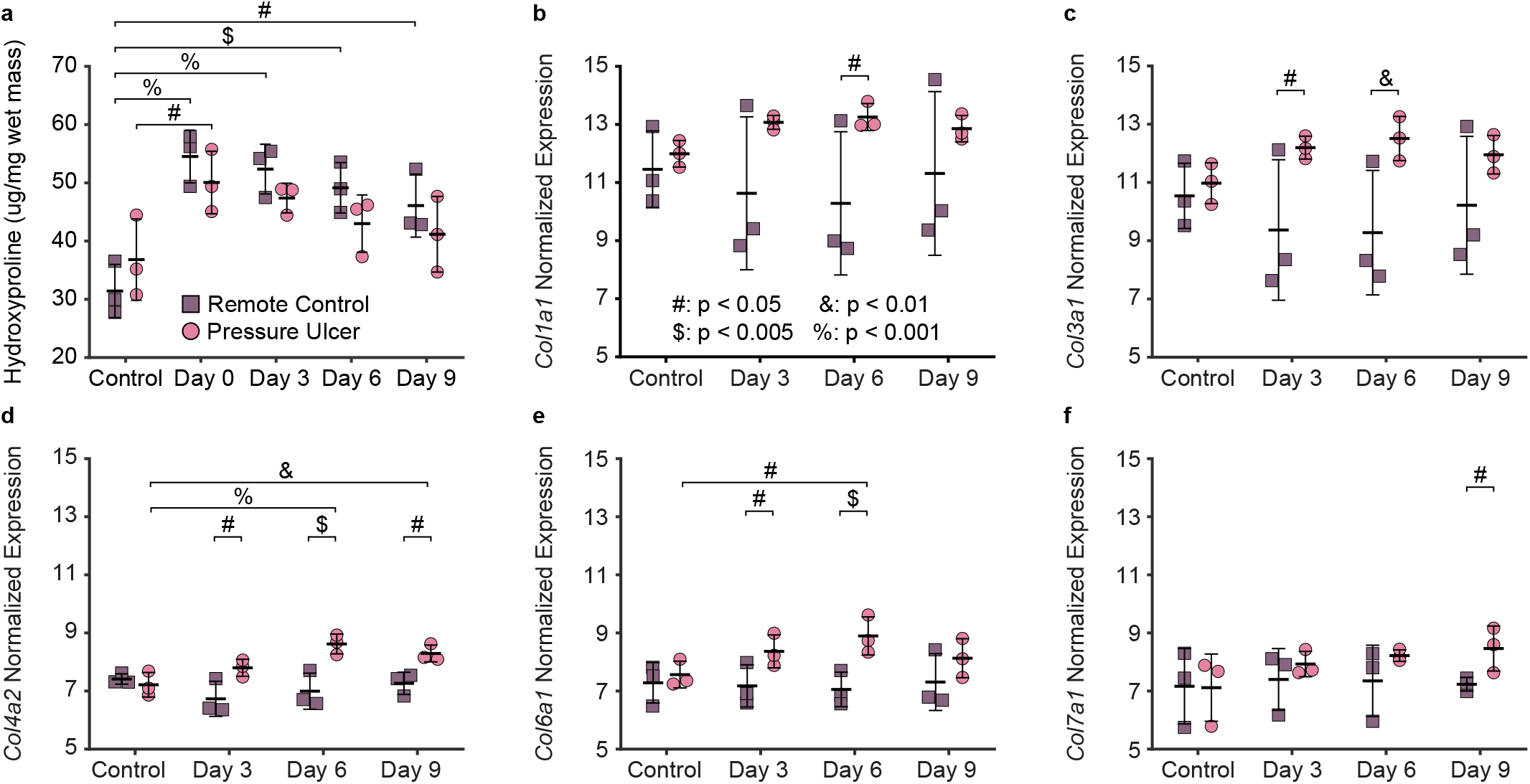
: Collagen remodeling during pressure ulcer progression at the protein and gene levels. **a**) Hydroxyproline-based quantification of total collagen content (*µ*g/mg wet mass) in remote control (RC) and pressure ulcer (PU) samples across time; collagen content increases at Day 0 in both RC and PU skin and then gradually decreases through Day 9. **b-f**) Normalized RNA-seq expression of specific collagen-encoding genes, including Col1a1, Col3a1, Col4a2, Col6a1, and Col7a1 in RC and PU samples across time; PU tissue shows time-dependent upregulation of fibrillar and nonfibrillar collagens, whereas RC skin displays no consistent increase

### Pressure ulcer formation triggers a pronounced inflammatory surge that peaks around Day 3 and gradually resolves

We next quantified inflammatory markers at both the protein and gene levels (Figure 7). ELISA measurements revealed consistently low cytokine levels in RC samples, whereas PU samples exhibited strong, marker-specific increases over time (Figure 7**a–f**). MCP-1, VEGF, and TGF-*β*1 increased as early as Day 0 and remained elevated through Days 3-9. In contrast, MPO, TNF-*α*, and IL-1*β* rose more prominently after reperfusion, peaking around Day 3 and remaining elevated at Days 6 and 9. RNA-seq analysis (Control and Days 3, 6, and 9) demonstrated time-dependent gene expression consistent with this post-reperfusion response (Figure 7**g–l**). *Il1b* showed marked upregulation in pressure ulcers at Day 3 and remained elevated, although expression slightly decreased over time. *Ccl2* and *Tgfb1* similarly increased in ulcer tissue at Day 3 but declined by Day 9. In contrast, *Tnf* exhibited comparatively modest gene-level changes, while *Vegfa* displayed smaller but detectable increases, most apparent at Day 3. We also observed increased expression of *Mmp8*, a matrix metalloproteinase that can contribute to extracellular matrix turnover, indicating that matrix-regulating genes are modulated alongside inflammatory mediators in this model. Overall, these protein and gene expression changes demonstrate a rapid inflammatory response initiated early after ischemia, followed by a pronounced post-reperfusion inflammatory peak and partial resolution by Day 9.

**Figure 7:**
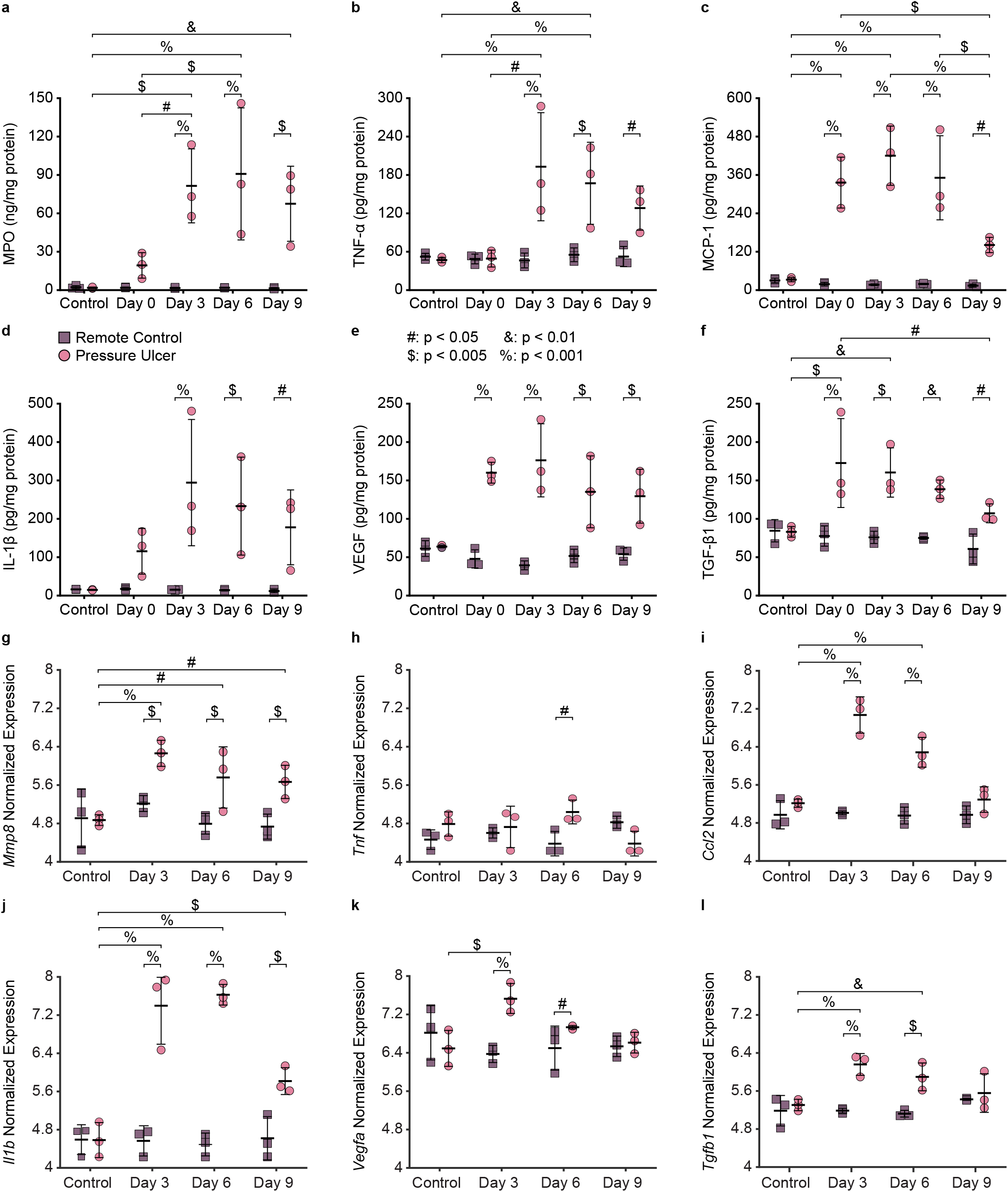
: Pressure ulcers exhibit a strong and sustained inflammatory response following ischemia-reperfusion injury. **a-f**) ELISA quantification of MPO, TNF-*α*, MCP-1, IL-1*β*, VEGF, and TGF-*β*1 in remote control (RC) and pressure ulcer (PU) samples over time; PU samples show cytokine levels that rise rapidly after injury, peak around Day 3, and remain elevated relative to RC through Day 9. **g-l**) Normalized RNA-seq expression of genes encoding the corresponding inflammatory and angiogenic mediators (*Mmp8, Tnf, Ccl2, Il1b, Vegfa, Tgfb1*); PU tissues display significantly higher gene levels than RC, consistent with the protein measurements

## Discussion

In this study, we addressed a critical gap in pressure ulcer research: the lack of longitudinal, quantitative data linking tissue mechanics, structural remodeling, inflammation, and gene-level responses. By integrating inverse finite element modeling, spatial histology, cytokine profiling, collagen quantification, and RNA-seq, we provide a comprehensive characterization of the spatiotemporal mechanical and biological co-evolution of pressure ulcers. Our findings demonstrate that pressure ulcer progression involves coordinated structural, mechanical, inflammatory, and collagen-associated extracellular matrix remodeling that evolves over distinct temporal scales. Together, these multi-scale, multi-modal data provide a quantitative foundation for developing predictive mechanobiological models of pressure ulcer progression and healing [13].

Pressure ulcer formation at Day 0 is associated with changes in matrix and fiber mechanics. Ulcer regions in pressure ulcer (PU) samples exhibit higher matrix moduli than surrounding skin at Day 0, followed by a gradual decrease over time, demonstrating continued evolution of tissue mechanical behavior throughout ulcer progression. Similar mechanical heterogeneity has been reported in acute excisional wounds, where local mechanical measurements and model-based fitting reveal a stiff wound core with a larger non-collagenous contribution than surrounding skin [42, 43]. In the skin immediately surrounding ulcers, the fiber modulus *k*_1_ decreases at Day 0. Additionally, the fiber stiffening parameter *k*_2_ is elevated at Day 0, decreases by Day 3, and remains lower than the control thereafter. These findings demonstrate that changes in the mechanical properties of the skin extend beyond the ulcer region.

Collagen remodeling follows a distinct temporal pattern. Total collagen content, measured as hydroxyproline per unit wet mass, increases early during ulcer progression in both PU and remote control (RC) samples, whereas transcriptional upregulation of collagen-related genes is more pronounced within PU samples at later time points. The early elevation in collagen content, particularly at Day 0 right after ischemia, likely reflects pressure-induced compaction of the tissues and changes in the balance between collagen synthesis and degradation rather than a simple net gain of stable collagen [53, 45]. In contrast, the subsequent upregulation of collagen-related genes in PU samples at Days 3 and 6 is consistent with the activation of extracellular matrix synthesis during the reparative phase, when fibroblasts and myofibroblasts synthesize new collagen in response to TGF-*β* signaling among other cytokines [34, 45]. Prior work in chronically ischemic but non-ulcerated human skin has shown that hypoxia can increase collagen synthesis and matrix metalloproteinase-mediated degradation, and can weaken dermal mechanical integrity by promoting collagen that is less stable and readily degradable, underscoring that increases in collagen synthesis or bulk collagen content do not necessarily translate into stronger or more mechanically resilient tissue under ischemic conditions [6, 7].

Interestingly, total collagen content also increases in RC samples without a corresponding transcriptional response. This divergence suggests that bulk collagen content and gene expression follow distinct temporal dynamics. In RC samples, elevated collagen content could arise from indirect responses to ischemia and reperfusion, such as changes in tissue hydration, tissue compaction, and matrix degradation kinetics [6, 1, 19]. Because we did not directly measure collagen crosslinking, fiber organization, non-collagenous matrix components, or water content, we cannot definitively assign mechanisms; however, our data collectively suggest that pressure ulcer progression involves early collagen densification and later, collagen synthesis within ulcer regions, and that ischemia-reperfusion-induced collagen remodeling extends into remote skin regions.

Together, the mechanical and collagen measurements provide complementary perspectives on tissue remodeling during the progression of pressure ulcers. Bulk collagen quantification characterizes changes in collagen abundance, but does not fully capture the evolving mechanical behavior of the tissue. Conversely, mechanical testing provides functional information that cannot be inferred from collagen content alone. Integrating these complementary measurements reveals that mechanical evolution and extracellular matrix remodeling follow distinct but overlapping temporal patterns, providing a more comprehensive understanding of pressure ulcer progression. Importantly, the mechanical characterization is enabled by our inverse finite element framework, which incorporates experimentally measured spatial thickness heterogeneity to account for geometric variability across samples. By fitting both force-displacement data and full-field displacement measurements, the framework strengthens parameter identification and enhances the physiological relevance of the estimated matrix and fiber material properties, in line with recent work emphasizing the impact of thickness and structural heterogeneity on soft tissue biomechanics [26, 28].

Furthermore, the inflammatory response in this model follows a dynamic temporal pattern. PU samples show elevated MPO, TNF-*α*, MCP-1, IL-1*β*, VEGF, and TGF-*β*1 relative to control tissues, with chemokines and growth factors such as MCP-1, VEGF, and TGF-*β*1 already increased at Day 0 and remaining high through Days 3-9, whereas neutrophil-associated MPO and classical pro-inflammatory cytokines TNF-*α* and IL-1*β* rise more prominently after reperfusion, peak around Day 3, and stay above control levels at later time points. In murine skin wound models, TNF-*α* and IL-1*β* rise rapidly after injury, exhibit a peak around 72 hours, and remain elevated during the early repair phase before declining substantially over the subsequent 1-2 weeks as wounds transition into proliferative and remodeling phases [21, 3, 23]. Consistent with our findings, compression-based pressure-ulcer models report sustained upregulation of inflammatory cytokines over the first several days after injury, including TNF-*α* and IL-1*β* and other inflammatory mediators [24]. RNA-seq analysis in our study further supports upregulation of corresponding gene expression during this phase, indicating a transcriptionally active inflammatory and angiogenic response in PU samples that decreases between Days 3 and 9 but does not fully return to baseline over this time window, suggesting incomplete resolution of inflammation over the 9-day period. This prolonged inflammatory response is consistent with previous experimental and clinical studies demonstrating that pressure ulcers heal slower than uncomplicated acute excisional wounds, with persistent inflammation, delayed vascular regeneration, and prolonged tissue remodeling [16, 33, 14, 41].

Complementing these bulk inflammatory measurements, spatial histology showed early suppression of vascular and macrophage markers in the ulcer centers followed by sustained, region-dependent increases during pressure ulcer progression. CD31^+^ endothelial staining was reduced within ulcer centers at Day 0, consistent with ischemic vascular compromise, but became markedly elevated in the side and deep regions at Days 3-6 and remained higher in ulcer centers at Day 9, indicating persistent neovascular growth and redistribution of vessels across the ulcer bed. F4/80^+^ macrophage signal displayed a similar response. Interestingly, macrophage staining in RC samples decreased at Day 3 while side and deep regions of PU samples exhibited strong increases, demonstrating that macrophages were recruited toward the ulcer site. F4/80^+^ staining remained elevated at Day 9 with increased signals in the ulcer centers. Together, our cytokine and gene-expression measurements and spatial CD31^+^ and F4/80^+^ maps show the temporal evolution of inflammatory and vascular remodeling in our pressure-ulcer model and provide a quantitative dataset that can support future mechanistic and computational models of pressure-ulcer pathophysiology. Future work with spatially resolved cytokine profiling could further link local VEGF and chemokine gradients to these CD31^+^ and F4/80^+^ patterns.

Despite the strengths of our work, this study has limitations. We did not directly measure collagen crosslink density, anisotropy (fiber orientation distribution), or other microstructural features (e.g. fiber diameter, waviness distributions), which limits our ability to assign specific structural mechanisms to the observed changes in mechanical parameters. Future studies could couple planar biaxial testing with in-situ multiphoton or confocal imaging to relate collagen fiber architecture to regional material properties, using biaxial devices specifically designed for image-based mechanical studies [32]. We also did not perform RNA-seq at Day 0 due to limited tissue mass and a lack of viable cells within the ulcer region, preventing direct evaluation of immediate transcriptional responses following ischemia. Our RNA-seq analysis was performed on bulk tissue samples, whereas sterile inflammation following ischemia reperfusion, ulcer formation, angiogenesis, and extracellular matrix remodeling involve diverse cell types (including keratinocytes, fibroblasts, endothelial cells, neutrophils, and macrophages), which interact through complex signaling cascades. The bulk RNA-seq data sheds light on some of the underlying biological mechanisms at play, but single-cell RNA-seq or spatial transcriptomics would be needed to reconstruct the role of each cell type in this complex process [4, 29]. Although our bulk RNA-seq dataset contains additional information on matrix-regulating genes, including other metalloproteinases and tissue inhibitors of metalloproteinases, in this study we focused on collagen-encoding genes and cytokines that correspond to our protein measurements; the full dataset will be made available to support future analyses that more deeply interrogate extracellular matrix turnover and its regulation during pressure ulcer progression.

Overall, our work establishes a multi-scale, multi-modal quantitative foundation linking mechanical evolution, inflammatory activation, and collagen-associated extracellular matrix remodeling in pressure ulcer progression. By capturing tissue geometry, mechanics, cellular activity, and gene expression within the same longitudinal framework, we provide data directly applicable to predictive mechanobiological models of pressure ulcer formation and healing. Incorporating spatiotemporal mechanical properties and biological responses into such models will be essential to improve their predictive capability and advance therapeutic strategies.

## Data Availability

The trimmed fastq RNA sequencing files will be available in the National Center for Biotechnology Information Sequence Read Archive. All the other data will be uploaded to our Data Repository and become available upon publication.

## Author Contributions

**Chien-Yu Lin:** Conceptualization, Methodology, Software, Formal Analysis, Investigation, Data Curation, Writing – Original Draft, Writing – Review & Editing, Visualization. **Shreya Sreedhar:** Formal Analysis, Investigation, Data Curation, Writing – Original Draft, Writing – Review & Editing, Visualization. **Matthew J. Lohr:** Software, Formal Analysis, Writing – Original Draft, Writing – Review & Editing, Visualization. **Colton J. Kostelnik:** Investigation, Writing – Review & Editing. **Alberto Madariaga:** Investigation, Writing – Review & Editing. **Adrian B. Tepole:** Conceptualization, Writing – Review & Editing, Supervision, Funding acquisition. **Manuel K. Rausch:** Conceptualization, Methodology, Writing – Original Draft, Writing – Review & Editing, Supervision, Funding acquisition.

## Acknowledgments

This work was supported by NSF grants to MKR (1916663) and ABT (1916668), an AHA Predoctoral Fellowship to CYL (25PRE1363276), an NIH T32 Training Fellowship to SS (T32 EB007507), an NIH Predoctoral Fellowship to MJL (F31HL176137), and an AHA Postdoctoral Fellowship to CJK (25POST1375865). The opinions, findings, and conclusions, or recommendations expressed are those of the authors and do not necessarily reflect the views of the National Science Foundation and the National Heart, Lung, and Blood Institute of the National Institutes of Health.

## References

[1] Beatty, R., Farina, P.P., O’Dwyer, J., Dockery, P., Stanley, A., 2025. Understanding the impact of reperfusion in the development of a safe compression therapy. Microscopy and Microanalysis: The Official Journal of Microscopy Society of America, Microbeam Analysis Society, Microscopical Society of Canada 31, ozaf056. doi:10.1093/mam/ozaf056.

[2] Blaber, J., Adair, B., Antoniou, A., 2015. Ncorr: Open-source 2d digital image correlation matlab software. Experimental Mechanics 55, 1105–1122. doi:10.1007/s11340-015-0009-1.

[3] Cañedo-Dorantes, L., Cañedo-Ayala, M., 2019. Skin acute wound healing: A comprehensive review. International Journal of Inflammation 2019, 3706315. doi:10.1155/2019/3706315.

[4] Cheng, X.C., Tong, W.Z., Rui, W., Feng, Z., Shuai, H., Zhe, W., 2024. Single-cell sequencing technology in skin wound healing. Burns & Trauma 12, tkae043. doi:10.1093/burnst/tkae043.

[5] Coleman, S., Nixon, J., Keen, J., Wilson, L., McGinnis, E., Dealey, C., Stubbs, N., Farrin, A., Dowding, D., Schols, J.M., Cuddigan, J., Berlowitz, D., Jude, E., Vowden, P., Schoonhoven, L., Bader, D.L., Gefen, A., Oomens, C.W., Nelson, E.A., 2014. A new pressure ulcer conceptual framework. Journal of Advanced Nursing 70, 2222–2234. doi:10.1111/jan.12405.

[6] Dalton, S.J., Mitchell, D.C., Whiting, C.V., Tarlton, J.F., 2005. Abnormal extracellular matrix metabolism in chronically ischemic skin: A mechanism for dermal failure in leg ulcers. Journal of Investigative Dermatology 125, 373–379. doi:10.1111/j.0022-202X.2005.23789.x.

[7] Dalton, S.J., Whiting, C.V., Bailey, J.R., Mitchell, D.C., Tarlton, J.F., 2007. Mechanisms of chronic skin ulceration linking lactate, transforming growth factor-β, vascular endothelial growth factor, collagen remodeling, collagen stability, and defective angiogenesis. Journal of Investigative Dermatology 127, 958–968. doi:10.1038/sj.jid.5700651.

[8] Demarré, L., Van Lancker, A., Van Hecke, A., Verhaeghe, S., Grypdonck, M., Lemey, J., Annemans, L., Beeckman, D., 2015. The cost of prevention and treatment of pressure ulcers: A systematic review. International Journal of Nursing Studies 52, 1754–1774. doi:10.1016/j.ijnurstu.2015.06.006.

[9] Gasser, T.C., Ogden, R.W., Holzapfel, G.A., 2006. Hyperelastic modelling of arterial layers with distributed collagen fibre orientations. Journal of the Royal Society, Interface 3, 15–35. doi:10.1098/rsif.2005.0073.

[10] Gefen, A., 2007. The biomechanics of sitting-acquired pressure ulcers in patients with spinal cord injury or lesions. International Wound Journal 4, 222–231. doi:10.1111/j.1742-481X.2007.00330.x.

[11] Gefen, A., Brienza, D.M., Cuddigan, J., Haesler, E., Kottner, J., 2021. Our contemporary understanding of the aetiology of pressure ulcers/pressure injuries. International Wound Journal 19, 692–704. doi:10.1111/iwj.13667.

[12] Gordon, S., Roberti, A., Yona, S., Lin, H.H., 2025. F4/80, the plasma membrane antigen of mouse macrophages, an historic journey. Journal of Leukocyte Biology 117. doi:10.1093/jleuko/qiaf126.

[13] Guo, Y., Mofrad, M.R.K., Tepole, A.B., 2022. On modeling the multiscale mechanobiology of soft tissues: Challenges and progress. Biophysics Reviews 3, 031303. doi:10.1063/5.0085025.

[14] Jiang, L., Dai, Y., Cui, F., Pan, Y., Zhang, H., Xiao, J., Xiaobing, F.U., 2014. Expression of cytokines, growth factors and apoptosis-related signal molecules in chronic pressure ulcer wounds healing. Spinal Cord 52, 145–151. doi:10.1038/sc.2013.132.

[15] Jin, T., Fu, Z., Zhou, L., Chen, L., Wang, J., Wang, L., Yan, S., Li, T., Jin, P., 2024. GelMA loaded with platelet lysate promotes skin regeneration and angiogenesis in pressure ulcers by activating STAT3. Scientific Reports 14, 18345. doi:10.1038/s41598-024-67304-2.

[16] Kasuya, A., Sakabe, J.i., Tokura, Y., 2014. Potential application of in vivo imaging of impaired lymphatic duct to evaluate the severity of pressure ulcer in mouse model. Scientific Reports 4, 4173. doi:10.1038/srep04173.

[17] Keenan, B.E., Evans, S.L., Oomens, C.W.J., 2022. A review of foot finite element modelling for pressure ulcer prevention in bedrest: Current perspectives and future recommendations. Journal of Tissue Viability 31, 73–83. doi:10.1016/j.jtv.2021.06.004.

[18] Khan, A.A., Alsahli, M.A., Rahmani, A.H., 2018. Myeloperoxidase as an active disease biomarker: Recent biochemical and pathological perspectives. Medical Sciences 6, 33. doi:10.3390/medsci6020033.

[19] Khorami, F., Foroutan, Y., Sparrey, C.J., 2025. Investigating the impact of dehydration and hydration on In-Vivo hip soft tissue biomechanics. PLOS One 20, e0328054. doi:10.1371/journal.pone.0328054.

[20] Koh, T.J., DiPietro, L.A., 2011. Inflammation and wound healing: the role of the macrophage. Expert Reviews in Molecular Medicine 13, e23. doi:10.1017/S1462399411001943.

[21] Kondo, T., Ohshima, T., 1996. The dynamics of inflammatory cytokines in the healing process of mouse skin wound: A preliminary study for possible wound age determination. International Journal of Legal Medicine 108, 231–236. doi:10.1007/BF01369816.

[22] Kostelnik, C.J., Meador, W.D., Lin, C.Y., Mathur, M., Malinowski, M., Jazwiec, T., Malinowska, Z., Piekarska, M.L., Gaweda, B., Timek, T.A., Rausch, M.K., 2025. Tricuspid valve leaflet remodeling in sheep with biventricular heart failure: A comparison between leaflets. Acta Biomaterialia 198, 234–244. doi:10.1016/j.actbio.2025.03.052.

[23] Kuninaka, Y., Ishida, Y., Ishigami, A., Nosaka, M., Matsuki, J., Yasuda, H., Kofuna, A., Kimura, A., Furukawa, F., Kondo, T., 2022. Macrophage polarity and wound age determination. Scientific Reports 12, 20327. doi:10.1038/s41598-022-24577-9.

[24] Kurose, T., Hashimoto, M., Ozawa, J., Kawamata, S., 2015. Analysis of gene expression in experimental pressure ulcers in the rat with special reference to inflammatory cytokines. PLOS ONE 10, e0132622. doi:10.1371/journal.pone.0132622.

[25] Limbert, G., 2017. Mathematical and computational modelling of skin biophysics: a review. Proceedings. Mathematical, Physical, and Engineering Sciences 473, 20170257. doi:10.1098/rspa.2017.0257.

[26] Lin, C.Y., Mathur, M., Malinowski, M., Timek, T.A., Rausch, M.K., 2023. The impact of thickness heterogeneity on soft tissue biomechanics: a novel measurement technique and a demonstration on heart valve tissue. Biomechanics and Modeling in Mechanobiology 22, 1487–1498. doi:10.1007/s10237-022-01640-y.

[27] Lin, C.Y., Sugerman, G.P., Kakaletsis, S., Meador, W.D., Buganza, A.T., Rausch, M.K., 2024. Sex- and age-dependent skin mechanics—A detailed look in mice. Acta Biomaterialia 175, 106–113. doi:10.1016/j.actbio.2023.11.032.

[28] Lipsky, Z.W., German, G.K., 2021. The precision of macroscale mechanical measurements is limited by the inherent structural heterogeneity of human stratum corneum. Acta Biomaterialia 130, 308–316. doi:10.1016/j.actbio.2021.05.035.

[29] Liu, Z., Bian, X., Luo, L., Björklund, Å.K., Li, L., Zhang, L., Chen, Y., Guo, L., Gao, J., Cao, C., Wang, J., He, W., Xiao, Y., Zhu, L., Annusver, K., Gopee, N.H., Basurto-Lozada, D., Horsfall, D., Bennett, C.L., Kasper, M., Haniffa, M., Sommar, P., Li, D., Landén, N.X., 2025. Spatiotemporal single-cell roadmap of human skin wound healing. Cell Stem Cell 32, 479–498.e8. doi:10.1016/j.stem.2024.11.013.

[30] Lohman, B.K., Weber, J.N., Bolnick, D.I., 2016. Evaluation of TagSeq, a reliable low-cost alternative for RNAseq. Molecular Ecology Resources 16, 1315–1321. doi:10.1111/1755-0998.12529.

[31] Lustig, A., Margi, R., Orlov, A., Orlova, D., Azaria, L., Gefen, A., 2021. The mechanobiology theory of the development of medical device-related pressure ulcers revealed through a cell-scale computational modeling framework. Biomechanics and Modeling in Mechanobiology 20, 851–860. doi:10.1007/s10237-021-01432-w.

[32] Madariaga, A., Lohr, M.J., Lin, C.Y., Fuhg, J., Buganza Tepole, A., Rausch, M., 2026. An affordable, openly-shared planar biaxial device to study the multi-scale mechanics of soft materials. Experimental Mechanics doi:10.1007/s11340-026-01267-5.

[33] Maldonado, A.A., Cristobal, L., Martín-López, J., Mallén, M., García-Honduvilla, N., Buján, J., 2014. A novel model of human skin pressure ulcers in mice. PLOS ONE 9, e109003. doi:10.1371/journal.pone.0109003.

[34] Mathew-Steiner, S.S., Roy, S., Sen, C.K., 2021. Collagen in wound healing. Bioengineering 8, 63. doi:10.3390/bioengineering8050063.

[35] Meador, W.D., Mathur, M., Sugerman, G.P., Malinowski, M., Jazwiec, T., Wang, X., Lacerda, C.M., Timek, T.A., Rausch, M.K., 2020a. The tricuspid valve also maladapts as shown in sheep with biventricular heart failure. eLife 9, e63855. doi:10.7554/eLife.63855.

[36] Meador, W.D., Sugerman, G.P., Story, H.M., Seifert, A.W., Bersi, M.R., Tepole, A.B., Rausch, M.K., 2020b. The regional-dependent biaxial behavior of young and aged mouse skin: A detailed histomechanical characterization, residual strain analysis, and constitutive model. Acta Biomaterialia 101, 403–413. doi:10.1016/j.actbio.2019.10.020.

[37] Mervis, J.S., Phillips, T.J., 2019. Pressure ulcers: Pathophysiology, epidemiology, risk factors, and presentation. Journal of the American Academy of Dermatology 81, 881–890. doi:10.1016/j.jaad.2018.12.069.

[38] Mondragon, N., Zito, P.M., 2025. Pressure injury, in: StatPearls. StatPearls Publishing, pp. 1–10.

[39] Murata, E., Fujii, J., 2025. Ischemia/reperfusion-associated oxidative stress is an aggravating factor for pressure ulcers. Journal of Clinical Biochemistry and Nutrition 76, 221–232. doi:10.3164/jcbn.25-28.

[40] Padula, W.V., Delarmente, B.A., 2019. The national cost of hospital-acquired pressure injuries in the united states. International Wound Journal 16, 634–640. doi:10.1111/iwj.13071.

[41] Palese, A., Luisa, S., Ilenia, P., Laquintana, D., Stinco, G., Di Giulio, P., PARI-ETLD Group, 2015. What is the healing time of Stage II pressure ulcers? Findings from a secondary analysis. Advances in Skin & Wound Care 28, 69–75. doi:10.1097/01.ASW.0000459964.49436.ce.

[42] Pensalfini, M., Haertel, E., Hopf, R., Wietecha, M., Werner, S., Mazza, E., 2018. The mechanical fingerprint of murine excisional wounds. Acta Biomaterialia 65, 226–236. doi:10.1016/j.actbio.2017.10.021.

[43] Pensalfini, M., Tepole, A.B., 2023. Mechano-biological and bio-mechanical pathways in cutaneous wound healing. PLOS Computational Biology 19, e1010902. doi:10.1371/journal.pcbi.1010902.

[44] Saito, Y., Hasegawa, M., Fujimoto, M., Matsushita, T., Horikawa, M., Takenaka, M., Ogawa, F., Sugama, J., Steeber, D.A., Sato, S., Takehara, K., 2008. The loss of MCP-1 attenuates cutaneous ischemia–reperfusion injury in a mouse model of pressure ulcer. Journal of Investigative Dermatology 128, 1838–1851. doi:10.1038/sj.jid.5701258.

[45] Singh, D., Rai, V., Agrawal, D.K., 2023. Regulation of collagen I and collagen III in tissue injury and regeneration. Cardiology and cardiovascular medicine 7, 5–16. doi:10.26502/fccm.92920302.

[46] Sree, V.D., Rausch, M.K., Tepole, A.B., 2019a. Linking microvascular collapse to tissue hypoxia in a multiscale model of pressure ulcer initiation. Biomechanics and Modeling in Mechanobiology 18, 1947–1964. doi:10.1007/s10237-019-01187-5.

[47] Sree, V.D., Rausch, M.K., Tepole, A.B., 2019b. Towards understanding pressure ulcer formation: Coupling an inflammation regulatory network to a tissue scale finite element model. Mechanics Research Communications 97, 80–88. doi:10.1016/j.mechrescom.2019.05.003.

[48] Stadler, I., Zhang, R.Y., Oskoui, P., Whittaker, M.B.S., Lanzafame, R.J., 2004. Development of a simple, noninvasive, clinically relevant model of pressure ulcers in the mouse. Journal of Investigative Surgery 17, 221–227. doi:10.1080/08941930490472046.

[49] Uchiyama, A., Yamada, K., Perera, B., Ogino, S., Yokoyama, Y., Takeuchi, Y., Ishikawa, O., Motegi, S.i., 2015. Protective effect of MFG-e8 after cutaneous ischemia–reperfusion injury. Journal of Investigative Dermatology 135, 1157–1165. doi:10.1038/jid.2014.515.

[50] Visconti, A.J., Sola, O.I., Raghavan, P.V., 2023. Pressure injuries: Prevention, evaluation, and management. American Family Physician 108, 166–174.

[51] Yamazaki, S., Sekiguchi, A., Uchiyama, A., Fujiwara, C., Inoue, Y., Yokoyama, Y., Ogino, S., Torii, R., Hosoi, M., Akai, R., Iwawaki, T., Ishikawa, O., Motegi, S.i., 2020. Apelin/APJ signaling suppresses the pressure ulcer formation in cutaneous ischemia-reperfusion injury mouse model. Scientific Reports 10, 1349. doi:10.1038/s41598-020-58452-2.

[52] Yoshikawa, T.T., Livesley, N.J., Chow, A.W., 2002. Infected pressure ulcers in elderly individuals. Clinical Infectious Diseases 35, 1390–1396. doi:10.1086/344059.

[53] Zhou, S., Salisbury, J., Preedy, V.R., Emery, P.W., 2013. Increased collagen synthesis rate during wound healing in muscle. PLoS ONE 8, e58324. doi:10.1371/journal.pone.0058324.

[54] Ziraldo, C., Solovyev, A., Allegretti, A., Krishnan, S., Henzel, M.K., Sowa, G.A., Brienza, D., An, G., Mi, Q., Vodovotz, Y., 2015. A computational, tissue-realistic model of pressure ulcer formation in individuals with spinal cord injury. PLoS Computational Biology 11, e1004309. doi:10.1371/journal.pcbi.1004309.

