## Supplemental Information for "The Mechanical and Biological Evolution of Pressure Ulcer Formation and Healing in Mice"

### 1. Inverse Finite Element Analysis

We show exemplary fits to displacement fields at the maximum applied strain. These fits correspond to the same samples used for the force and displacement data shown in Figure 3.

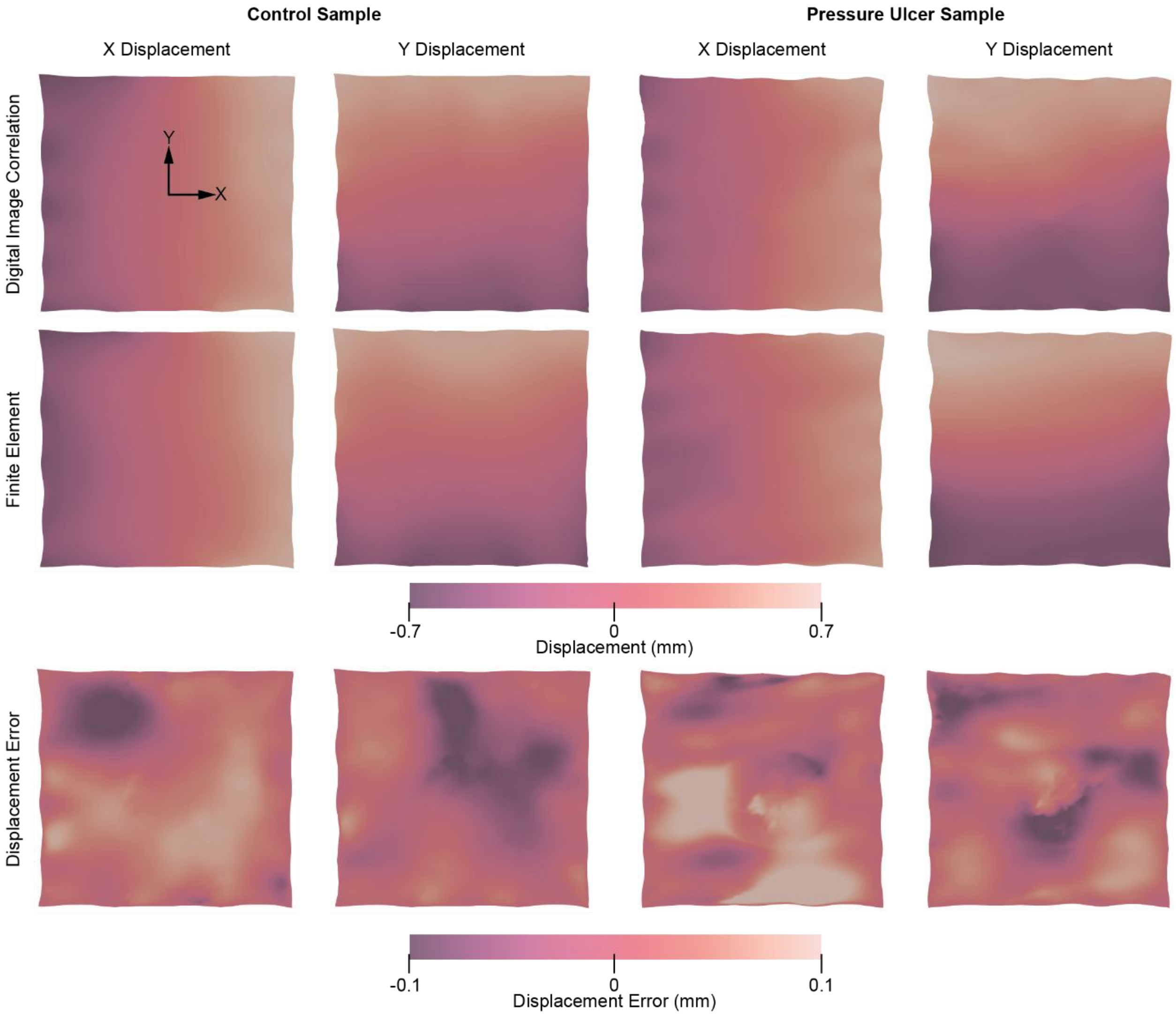

**Figure S1.** Displacement fields and errors for exemplary control (left) and pressure ulcer (right) samples at maximum applied strain. Experimentally measured displacement fields from digital image correlation are shown in the top row, model-predicted displacement fields using best-fit parameters in the middle row, and the corresponding error fields in the bottom row. Within each column, X (left) and Y (right) displacement components are shown, with a coordinate system indicating the X and Y directions. The normalized mean square errors (NMSEs) for the control sample are 0.985 (X displacement) and 0.984 (Y displacement), and for the pressure ulcer sample are 0.982 (X displacement) and 0.963 (Y displacement).

We used a least-squares approach to identify material parameters that minimized the error between measured and predicted forces and displacement fields. We set bounds on the parameters based on their admissible values except for the moduli $C_S$ and $C_U$, for which we set a lower bound of 1 Pa. Previous studies have shown that the softest regions of skin exhibit shear moduli on the order of 10 Pa; therefore, we restricted the moduli to values at least one order of magnitude below this estimate (Chi et al., 2024). We defined error vectors for each measurement. The force error vector was defined as

$$\boldsymbol{\varepsilon}_f = (1-w)(\boldsymbol{f}^E - \boldsymbol{f}^C), \tag{1}$$

where $\boldsymbol{f}^E$ are the measured forces, $\boldsymbol{f}^C$ are the computed forces, and $w \in (0,1)$ is a weight. The displacement error vector is defined as

$$\boldsymbol{\varepsilon}_d = w(\boldsymbol{d}^E - \boldsymbol{d}^C), \tag{2}$$

where $\boldsymbol{d}^E$ are the experimental displacements, and $\boldsymbol{d}^C$ are the computed displacements. We manually adjusted the weight to achieve high-quality fits to the data, as measured by the normalized mean squared error (NMSE), with higher values indicating better agreement between the model and experiment. We selected $w = 0.8$ for control samples and $w = 0.9$ for ulcer samples. We concatenated the error vectors and scaled the force error by $w_L = \mathrm{length}(\boldsymbol{d}^E)/\mathrm{length}(\boldsymbol{f}^E)$. To reduce the risk of convergence to a local minimum, we used 10 initial guesses with nonnegative NMSEs generated by Latin Hypercube Sampling and selected the solution that maximized the NMSE (i.e., best agreement).

Chi, W.-Y., Huang, H.-W., Lee, G., Cruz, C.J.G., Hughes, M.W., Tang, M., Shieh, S.-J., Yang, C.-C., 2024. Mechanical stiffness across skin layers in human: a pilot study. Tissue Barriers 13, 2437220. https://doi.org/10.1080/21688370.2024.2437220

**Table S1.** Average normalized mean squared error (NMSE) for all skin samples and ulcer samples. Shown are the total mean NMSE and the NMSE for each dataset used in the fitting process. The skin samples have higher NMSEs than the ulcer samples, and the fits to the force data are superior to those for the displacement fields, as demonstrated by mean NMSE values exceeding 0.9 for all sample types (with higher NMSE indicating better agreement).

| Sample Type | Mean | X-Displacement | Y-Displacement | X-Force | Y-Force |
|---|---|---|---|---|---|
| All | 0.93 | 0.90 | 0.92 | 0.95 | 0.96 |
| Skin | 0.96 | 0.95 | 0.96 | 0.96 | 0.96 |
| Ulcer | 0.90 | 0.83 | 0.88 | 0.95 | 0.96 |

### 2. Histology & Immunohistochemistry

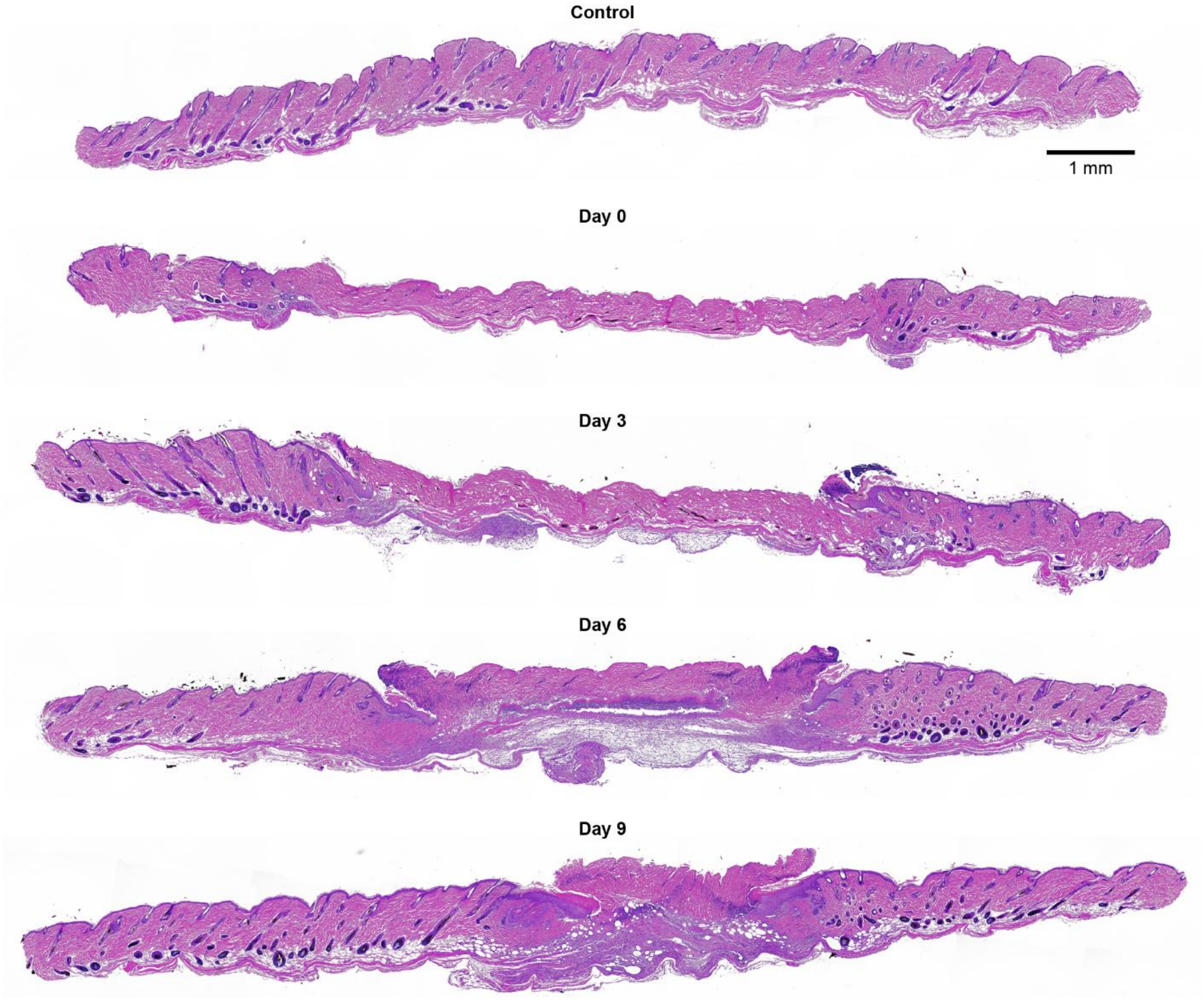

**Figure S2.** Representative H&E-stained sections from the pressure ulcer site show structural progression of tissue injury and healing. Sections are shown for control tissue and at Days 0, 3, 6, and 9 following ischemia-reperfusion injury. H&E staining highlights overall tissue architecture and the distribution of cell nuclei. Progressive disruption of dermal structure is observed at early time points, followed by gradual tissue reorganization during healing. Scale bar: 1 mm.

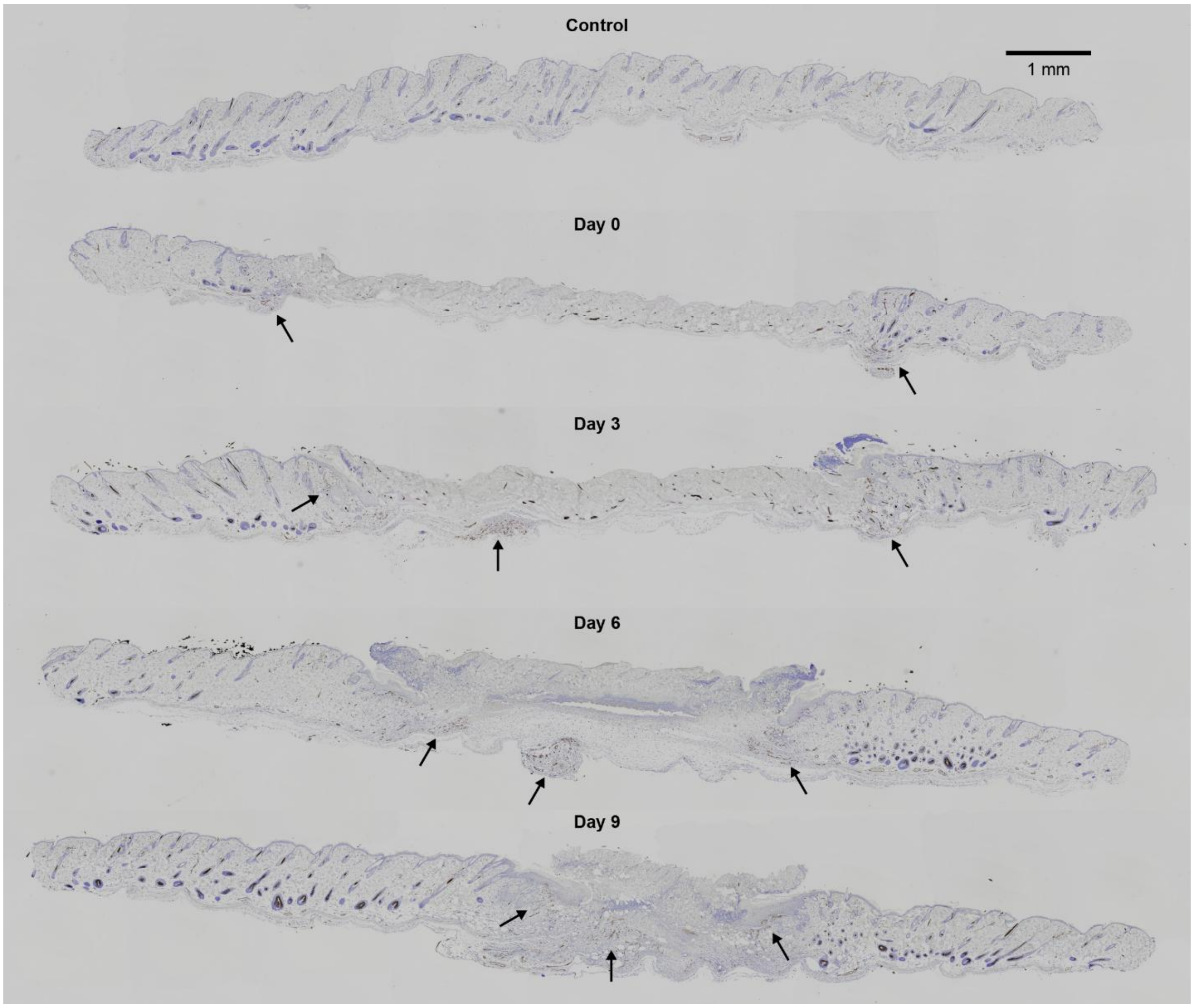

**Figure S3.** Representative CD31 immunohistochemistry sections from the pressure ulcer site show spatiotemporal changes in microvasculature. Sections are shown for control tissue and at Days 0, 3, 6, and 9 following ischemia-reperfusion injury. CD31+ staining identifies endothelial cells and vascular structures. Arrows indicate regions with increased CD31+ signal, with early suppression in the ulcer center followed by greater signal in peripheral and deeper regions during reperfusion. Scale bar: 1 mm.

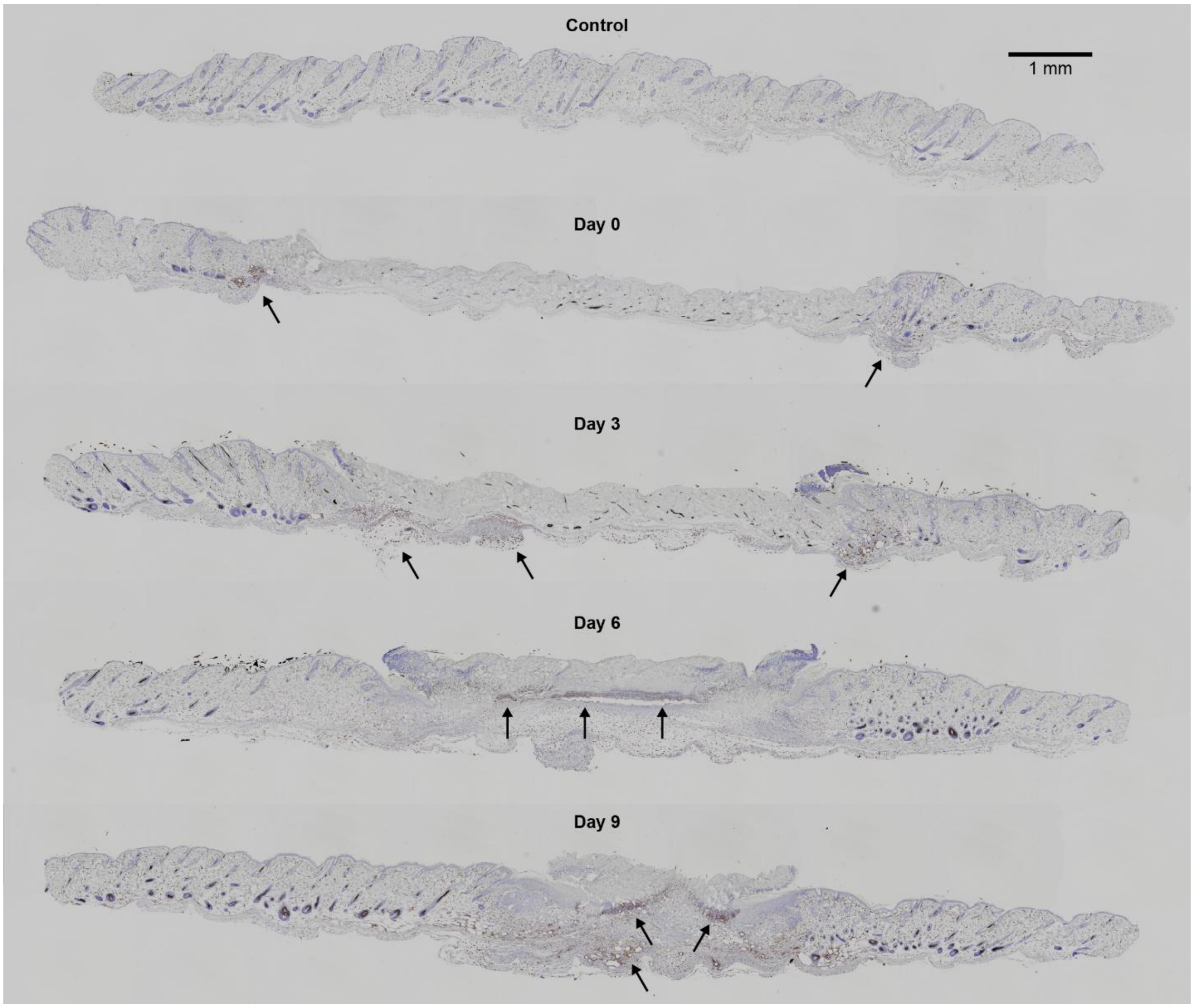

**Figure S4.** Representative F4/80 immunohistochemistry sections from the pressure ulcer site show spatiotemporal macrophage recruitment. Sections are shown for control tissue and at Days 0, 3, 6, and 9 following ischemia-reperfusion injury. F4/80+ staining identifies macrophages. Arrows indicate regions with increased F4/80+ signal, demonstrating initial depletion in the ulcer center followed by greater signal in peripheral and deeper regions during healing. Scale bar: 1 mm.

#### 3. RNA Sequencing

We show volcano plots to compare different conditions and highlight the extent of differential gene expression. Selected genes were then isolated and plotted across experimental groups and sample locations, as shown in Figures 6 and 7.

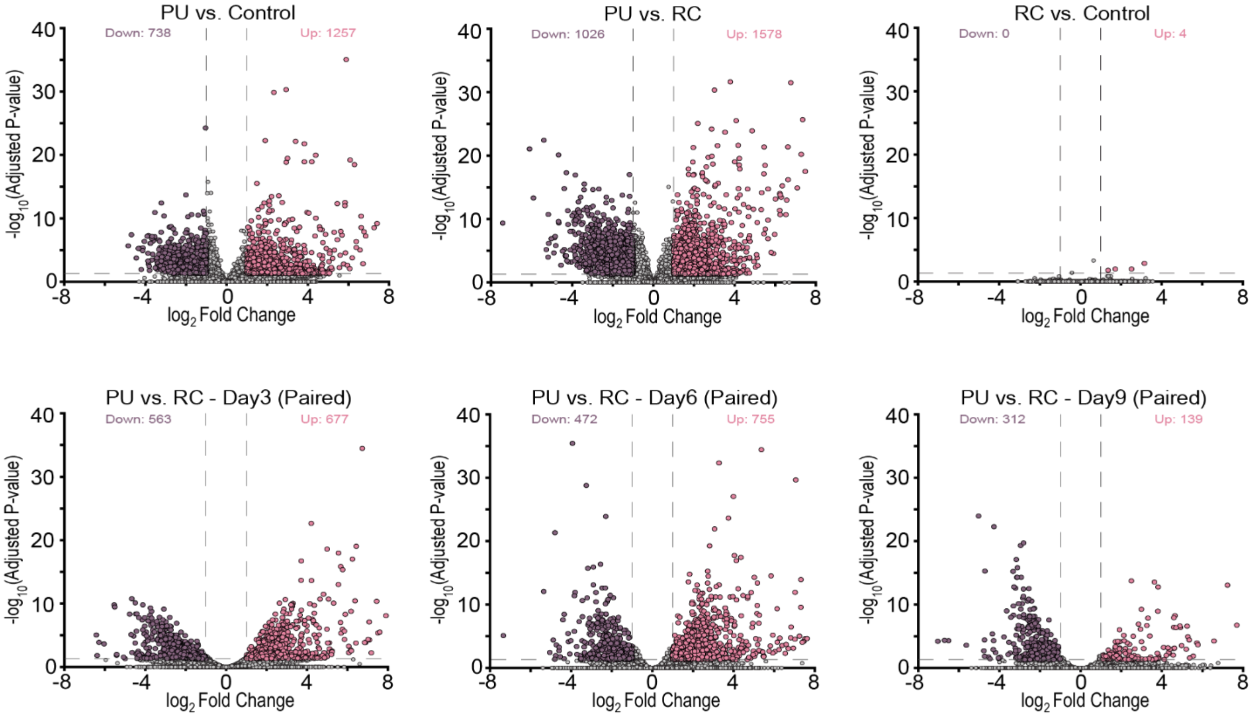

**Figure S5.** Volcano plots for samples from pressure ulcer (PU) sites vs. Control (left), PU vs. samples from remote control (RC) sites (middle), and RC vs. Control (right) are shown in the top row. Volcano plots for PU vs. RC for Days 3 (left), 6 (middle), and 9 (right) are shown in the bottom row. The volcano plots separated by day were paired by animal within each day. Differentially expressed genes (DEGs) that are significantly upregulated are highlighted in pink, and significantly downregulated DEGs are highlighted in purple. The number of significantly up- or downregulated genes is displayed in the top-right and top-left corners of each plot, respectively. A $\log_2$ (fold change) greater than 1 and a $-\log_{10}$ (adjusted p-value) greater than 0.05 were used as thresholds for determining significantly upregulated or downregulated genes.
